# Detection of genome-wide low-frequency mutations with Paired-End and Complementary Consensus Sequencing (PECC-Seq) revealed end-repair derived artifacts as residual errors

**DOI:** 10.1101/2019.12.22.886440

**Authors:** Xinyue You, Suresh Thiruppathi, Weiying Liu, Yiyi Cao, Mikihiko Naito, Chie Furihata, Masamitsu Honma, Yang Luan, Takayoshi Suzuki

**Author notes:** To whom correspondence should be addressed., Correspondence may also be addressed to Yang Luan.

## Abstract

To improve the accuracy and the cost-efficiency of next-generation sequencing in ultralow-frequency mutation detection, we developed the Paired-End and Complementary Consensus Sequencing (PECC-Seq), a PCR-free duplex consensus sequencing approach. PECC-Seq employed shear points as endogenous barcodes to identify consensus sequences from the overlap in the shortened, complementary DNA strands-derived paired-end reads for sequencing error correction. With the high accuracy of PECC-Seq, we identified the characteristic base substitution errors introduced by the end-repair process of mechanical fragmentation-based library preparations, which were prominent at the terminal 6 bp of the library fragments in the 5’-NpCpA-3’ or 5’-NpCpT-3’ trinucleotide context. As demonstrated at the human genome scale (TK6 cells), after removing these potential end-repair artifacts from the terminal 6 bp, PECC-Seq could reduce the sequencing error frequency to mid-10^−7^ with a relatively low sequencing depth. For TA base pairs, the background error rate could be suppressed to mid-10^−8^. In mutagen-treated TK6, slight increases in mutagen treatment-related mutant frequencies could be detected, indicating the potential of PECC-Seq in detecting genome-wide ultra-rare mutations. In addition, our finding on the patterns of end-repair artifacts may provide new insights in further reducing technical errors not only for PECC-Seq, but also for other next-generation sequencing techniques.

## INTRODUCTION

Next-generation sequencing (NGS) technologies have revolutionarily altered genomic explorations in the past decade. NGS makes genotyping of billions of bases simultaneously with relatively low cost possible and has been a routine in characterizing genomic variations in the fields of both basic research and clinical applications. However, as a trade-off for high throughput, accuracy of NGS is compromised compared with conventional techniques, with an error rate of approximately 10^−3^ to 10^−2^ (1–3). Nevertheless, in the scenario of clonal mutations, by sequencing sufficient homogeneous copies, random sequencing errors can be removed and reliable results can be obtained despite the current accuracy. By contrast, for heterogeneous samples, which are much more common in practice, the sensitivity of NGS techniques in detecting subclonal mutations is greatly limited by the error rates. Theoretically, the detection limit of the subclonal variants is determined by the accuracy of applied sequencing methods, which means approximately 1% with standard NGS accuracy (4).

With the increasing demand for the detection of low-frequency subclonal mutations in a variety of fields (e.g., cancer research, public health surveillance, metagenomics and forensics) (5–10), lowering the error rates and improving the sensitivity of the current NGS techniques have become an urgency. Besides optimizing the experimental conditions and bioinformatic algorithms, a third category, termed the molecular consensus sequencing strategy, has exhibited great potential in correcting sequencing artifacts (4). The principle for the consensus sequencing strategy is to get redundant copies that arise from same templates, which should be homogeneous, for error correction: true mutations should present in all or majority of the multiple copies, and any variant that only happens in part of the homogenous copies will be considered as sequencing artifacts. Generally, in order to get sufficient copies for consensus sequencing, PCR amplification is necessary. Besides, this strategy also requires molecular barcodes to identify the PCR-amplified copies that are generated from same starting templates. Based on this strategy, many attempts have been made to improve the accuracy of standard NGS techniques (4,11), e.g., Safe-Sequencing System (Safe-SeqS) (12), circle sequencing (13), bottleneck sequencing system (BotSeqS) (14), duplex sequencing (DupSeq) (15) and hypothesis alignment with weak overlap (Hawk-Seq^TM^) (16) for short-read platforms, as well as INC-seq (17) and circular consensus sequencing (CCS) (18) for long-read platforms. In short-read platforms, these consensus sequencing approaches have successfully improved the accuracy to a level of 10^−4^ to 10^−7^ or even lower (4,11).

Contrary to the high accuracy, due to the redundant nature of consensus sequencing strategies, the sequencing yield and cost become the major considerations, especially when applying such approaches to the genome-wide mutation screening. To create consensus sequences, for a single template, redundant PCR-amplified copies need to be sequenced for error correction; while theoretically, only one copy of the template is sequenced in conventional NGS techniques to avoid PCR duplicates. Thus, to obtain similar yields as conventional NGS methods, the raw sequencing depth required for consensus sequencing should be much higher, resulting in an exponential increase of sequencing cost. In addition, considering the biased nature of PCR amplification (19,20), more PCR cycles are sometimes required to get a uniform coverage for the loci that are difficult to amplify, which will further increase the average sequencing depth (4,11). The high sequencing depth makes these PCR amplification-based consensus sequencing approaches less cost-efficient when applied to large genomes and be more suitable for smaller genomic targets (1,4,11). Although BotSeqS has provided an excellent example in detecting genome-wide somatic mutations with a strategy of dilution before PCR amplification to reduce the sequencing depth (14), the sequencing efficiency is still very low and there is still scope for further improving the efficiency of consensus sequencing approaches.

To overcome the low sequencing depth-yield efficiency when applying consensus sequencing to high-accuracy low-frequency mutation detection at the whole genome scale, we developed a novel consensus sequencing method, the Paired-End and Complementary Consensus Sequencing (PECC-Seq). Differing from other consensus sequencing approaches, PECC-Seq employed a simple modified PCR-free sequencing workflow and attempted to increase the consensus-making efficiency (i.e., the number of reads utilized to create a consensus sequence (4)) of whole genome sequencing data by capitalizing the overlap in the shortened paired-end reads and information from complementary strands of the double-stranded DNA (dsDNA) rather than utilizing PCR-amplified copies for duplex consensus sequencing. With the shear points as endogenous tags, the library preparations and subsequent bioinformatic analysis were further simplified. In addition, we combined computational reduction of end-repair artifacts with PECC-Seq to further increase the accuracy. With a relatively low depth, PECC-Seq exhibited a high accuracy and the uniform coverage in the mutation detection of whole mammalian genomes.

## MATERIALS AND METHODS

### Cell culture and treatment

TK6 human lymphoblastoid cell line (TK6) was maintained in the RPMI-1640 medium containing 10% horse serum and treated with DMSO (the solvent control), 6 μg/mL methyl methanesulfonate (MMS, Sigma-Aldrich, MO, USA) or 12 μg/mL N-Nitroso-N-ethylurea (ENU, Sigma-Aldrich, MO, USA) for 4 h. The cells were then collected for the *TK* gene mutation assay following the protocol as previously described (21,22), and for the PECC-Seq analysis, respectively.

### Library preparation and sequencing

After treatment, genomic DNA of TK6 was extracted with the mtDNA extractor CT kit (WAKO, Osaka, Japan) and subjected to library preparation following the Illumina TruSeq DNA PCR-Free library preparation workflow with the prolonged ultrasonic shearing procedure to generate shorter double-stranded library fragments. Sequencing was performed on the Illumina HiSeqX platform at approximately 40 × sequencing depth with the read length of 2 × 150 bp.

Genomic DNA from the control TK6 was further subjected to the enzymatic fragmentation and library preparation with the Lotus DNA Library Prep Kit (Integrated Device Technology, CA, USA). Modified fragmentation time and AMPure XP beads (Beckman Coulter, CA, USA) selection procedure were applied to generate shorter library fragments for sequencing. The constructed library was then sequenced on the Illumina MiniSeq platform with a Mid Output Kit (Illumina, CA, USA).

### Data processing

Raw sequencing reads were subjected to adapter removal by using the Trimmomatic software with the palindrome mode (23) and then aligned to human reference genome hg38 with the Burrows-Wheeler aligner (BWA). After mapping, reads with multiple mappings were discarded and only properly mapped paired-end reads with mapping quality of 60 were retained. Complementary reads were extracted when the following criteria were met: 1) the two pairs of paired-end reads shared same mapping 5’ coordinates; 2) the two pairs of paired-end reads had different mapping orientations; and 3) no other paired-end reads had matched mapping 5’ coordinates. To generate consensus sequences from a pair of complementary paired-end reads, overlapped regions of which with 4 supporting bases (Figure 1C, the region within the red frame) were used and only positions with 4 identical bases were included for creating consensus bases.

**Figure 1.**
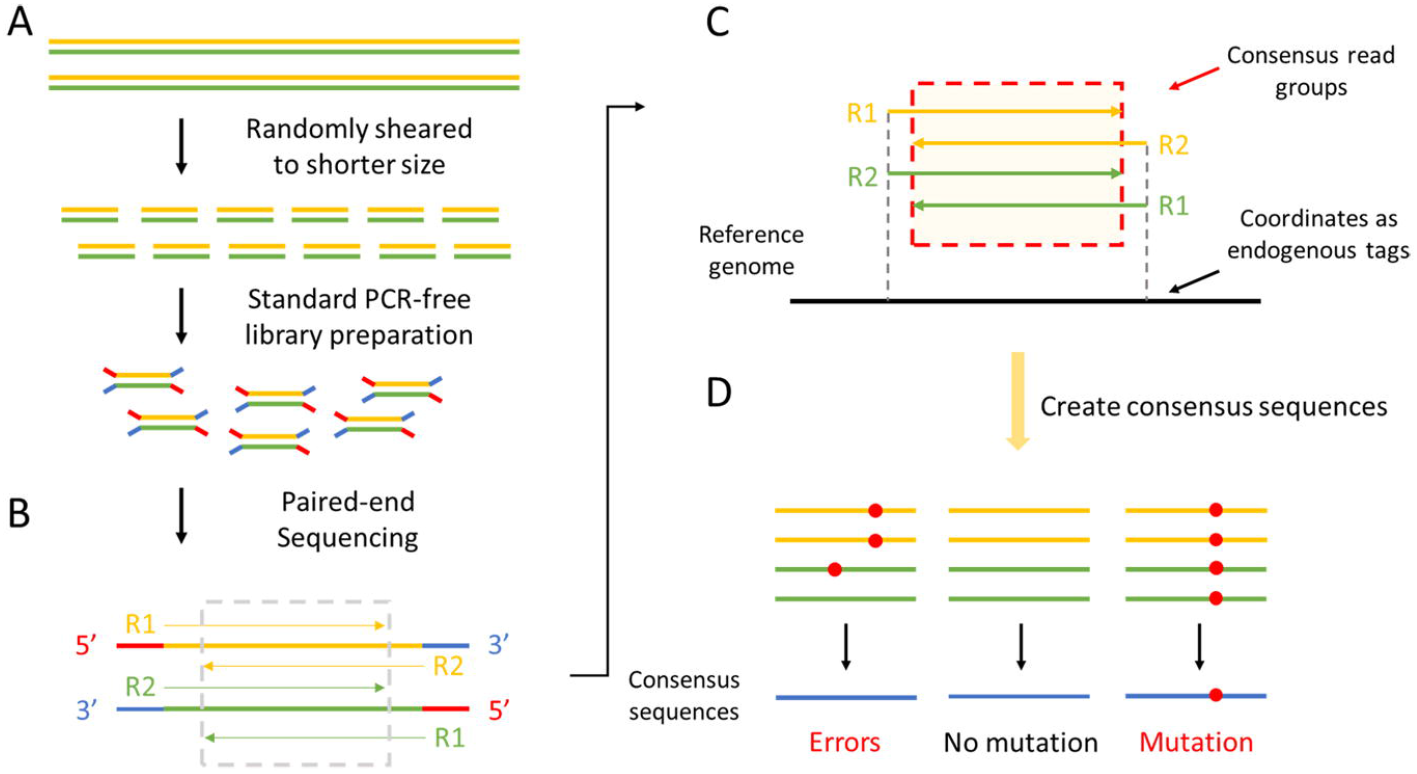
Overview of PECC-Seq. (**A**) Library preparation. A PCR-free library construction workflow was applied with prolonged ultrasonic shearing process to construct libraries with shortened fragment sizes. (**B**) Paired-end Sequencing. In the shortened library fragments, overlaps between the paired-end reads were obtained. (**C**) Reads processing. Mapping coordinates and mapping orientations served as endogenous tags to extract the consensus read groups. As the two templates arising from the DNA duplex were reverse complementary, the two pairs of reads shared same mapping coordinates (dashed lines) while the mapping orientations were opposite (the directions of R1 and R2). By utilizing the information in the overlap (the red frame), total 4 reads with identical information could be obtained to form duplex consensus sequences. (**D**) Creating concensus sequences and removing sequencing errors. At the given position, any variant that only occurred in part of the 4 bases was considered as an error and removed. Only positions with 4 identical bases would be included to make consensus sequences for further mutation analysis.

The resulting consensus bases were then filtered and candidate mutations were picked up with the following criteria: 1) variants happened solely in the consensus reads and not found in other reads of the variant sites; and 2) variants were not found in data of other treatment groups. All candidate mutants were further confirmed with the IGV browser. Further details on the data processing are provided in Supplementary Materials and Methods.

### Statistical Analysis

The differences between the mutant frequencies were evaluated with the *Poisson* test using the R software. A *p*-value of < 0.05 was considered statistically significant.

## RESULTS

To improve the consensus-making efficiency of the consensus sequencing strategy and enhance its practicability in large genomic targets, we designed the PECC-Seq, a PCR-free consensus sequencing approach (Figure 1). In this method, we modified the fragmentation protocol to construct the libraries with shortened fragment sizes (approximately 170 bp on average in the current work) (Figure 1A). Compared with standard sequencing libraries, this will ensure more overlaps between the read pairs after paired-end sequencing, in which the two paired-end reads share reverse complementary but identical information of the single-stranded templates (Figure 1B, as indicated in the grey frame). Owing to the double-stranded nature of DNA molecules, each sequenced single-stranded fragment has a reverse complementary strand that derived from the same individual dsDNA molecule. By capitalizing the overlap between paired-end reads and the complementary strands arising from dsDNA in combination, 4 reads sharing information of the same dsDNA templates (termed the consensus read group) can be obtained for creating duplex consensus sequences (Figure 1C, as indicated in the red frame). As the libraries were randomly sheared with ultrasonication and no PCR amplification was performed during library preparations, the probability of two individual fragments by chance to have same shear points was very low with the current sequencing depth (Supplementary Results). Thus, instead of exogenous barcodes, mapping 5’ coordinates (Figure 1C, as indicated with dashed lines) and mapping orientations (Figure 1C, as indicated by the directions of R1 and R2) of the paired-end reads were used as endogenous tags to label the reverse complementary templates-derived copies. At a given site, a true variant should present in all the four reads from the consensus read groups while sequencing errors and single strand damages could only be observed in part of the reads (Figure 1D).

### Apply PECC-Seq to whole genome sequencing

To evaluate the performance of PECC-Seq in genome-wide mutation detection, we applied it to TK6 human lymphoblastoid cell line (TK6) genome. After treatment with DMSO (the solvent control), or MMS and ENU, two known mutagens that widely serve as positive controls in gene mutation assays (22,24), the genomic DNA of TK6 was extracted and subjected to PECC-Seq analysis.

The number of obtained consensus bases and detected mutants are shown in Table 1, revealing mutant frequencies (MFs) of 0.76 × 10^−6^, 1.14 × 10^−6^ and 2.85 × 10^−6^ for control, MMS-treated and ENU-treated TK6, respectively. The observed indel frequencies were much lower by two orders than the frequencies of single nucleotide variants (SNVs). This suggests that PECC-Seq can efficiently lower the background error frequency from mid-10^−3^ to a level of 10^−7^ to 10^−6^ (Table S1). However, the resultant mutant frequencies are still higher than the background mutation frequencies determined in other studies (i.e., approximately 10^−8^ to 10^−7^) (25–27), indicating that there are some artifacts hidden in the detected mutants, especially for the observed SNVs.

**Table 1.** Number of mutants detected in treated TK6.

| Treatment | Consensus bases<br>($\times 10^8$) | Total<br>mutants | Total-MF<br>($\times 10^{-6}$) | SNV | SNV-MF<br>($\times 10^{-6}$) | Deletion | Insertion | Indel-MF<br>($\times 10^{-6}$) |
| --- | --- | --- | --- | --- | --- | --- | --- | --- |
| Control | 2.83 | 216 | 0.76 | 213 | 0.75 | 2 | 1 | 0.01 |
| MMS | 2.88 | 329 | 1.14 | 324 | 1.13 | 4 | 1 | 0.02 |
| ENU | 2.22 | 633 | 2.85 | 628 | 2.83 | 5 | 0 | 0.02 |

From the control, MMS-treated and ENU-treated TK6 libraries, 2.68 × 10^8^ bp (8.63 %), 2.72 ×10^8^ bp (8.77 %) and 2.12 × 10^8^ bp (6.85 %) of the genome could be covered, resulting in 1.06 ×, 1.06 × and 1.05 × consensus sequencing depth, respectively (Table S2). This indicates the scattered distribution of the consensus read groups and hence the uniform coverage of the genome can be achieved with PECC-Seq.

### Artifacts introduced during library preparations

Among the 213 SNVs obtained from the control TK6, we found that about half of the variants were aggregated in the terminal regions of the DNA templates. Same type of bias was also observed in chemical-treated groups, suggesting the error-prone end-repair process during library preparations may be implicated (4,28). To access this bias, we calculated the mutant frequencies by positions with given distances (bp) to the closest terminus of the DNA templates (i.e., the terminus of the library fragments, but not the end of the reads). As expected, the mutant frequencies in the distal regions were more than one order of magnitude higher than that of the interior regions (Figure 2), indicating bases in the terminal regions may suffer from the end-repair process and variants present in these sites were more likely to be artifacts rather than mutations. The error frequencies decreased significantly from the position of 7 bp to the terminus in the control data (*p* = 0.013) and ENU-treated data (*p* < 0.01). For MMS-treated data, the error frequency decreased significantly from the 8^th^ bp to the terminus (*p* < 0.01). Therefore, for the current data, we took the sites within 6 bp to the terminus as the distal region, and variants detected in this region were considered as the artifacts introduced during library preparation.

**Figure 2.**
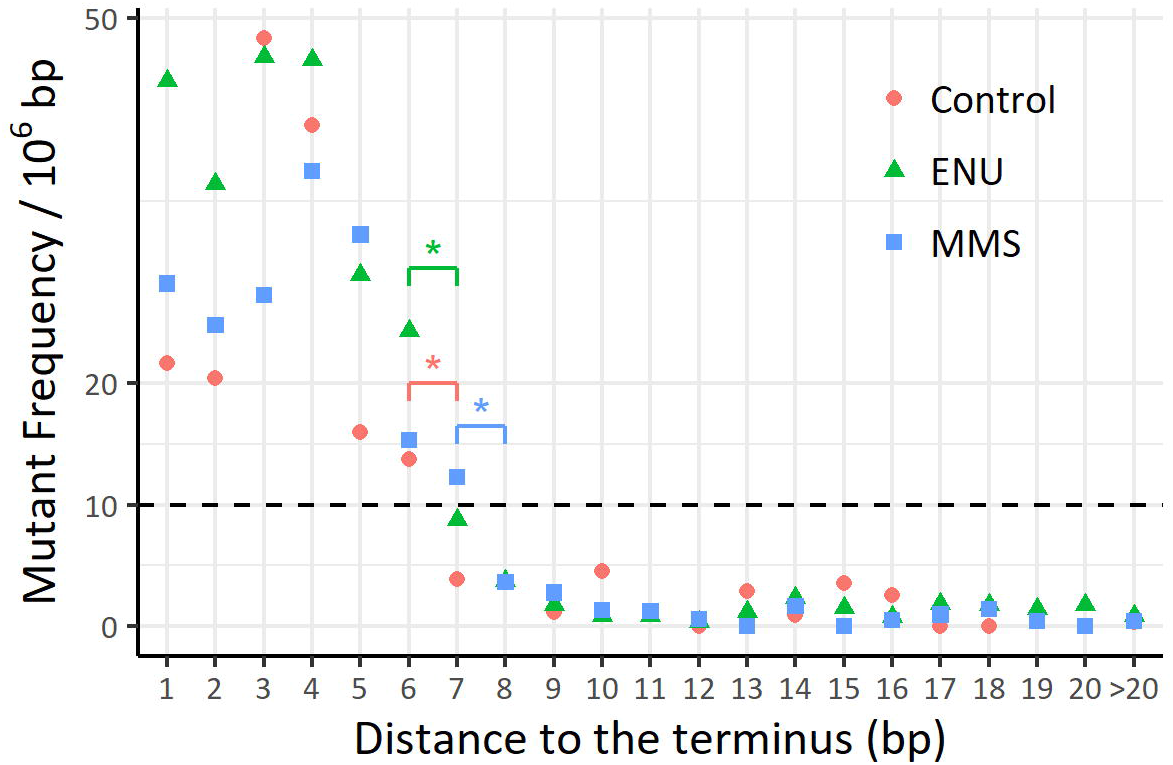
Mutant frequencies by positions with given distances to the terminus. The terminus is defined as the beginning or end of the DNA templates (library fragments). * *p* < 0.05.

The total 685 mutants found within the 6 bp-long distal regions (102 from control data, 180 from MMS-treated data and 403 from ENU-treated data) were subjected to further mutation spectrum construction with the 96-trinucleotide profile. These artifacts exhibited an obvious preference to the CG base pairs with 3’ flanked by A or T (i.e., 5’-NpCpA-3’ and 5’-NpCpT-3’) and displayed the highest error rate on CG > GC transversion, followed by CG > AT transversion and CG > TA transition (Figure 3, Figure S1).

**Figure 3.**
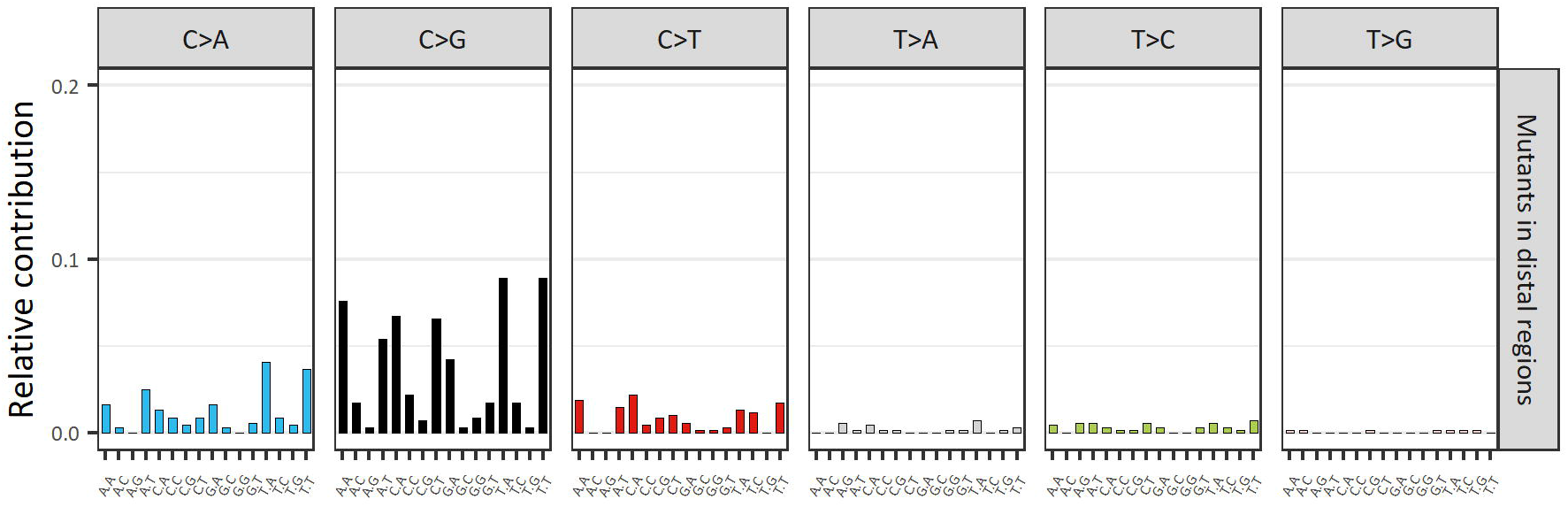
Profile of mutants detected in distal regions. The total 685 mutants obtained from the distal regions of the control data, MMS-treated data and ENU-treated data were merged for the 96-trinucleotide profile.

### Accuracy of PECC-Seq after removing artifacts in distal regions

Base on the distribution pattern of the terminal bias, we removed the error-prone 6 base pairs from distal regions to eliminate potential end-repair artifacts. After correcting the terminal bias, the accuracy of PECC-Seq was improved by about 2-fold, resulting in the mutant frequencies of 0.41 × 10^−6^ for the control, 0.53 × 10^−6^ for MMS-treated TK6 and 1.08 × 10^−6^ for ENU-treated TK6 (Table 2, Table S1). Although still slightly higher than the background mutation frequencies of human genomes, the resultant mutant frequency of the control data did exhibit a comparable level with that of human tissues determined by BotSeqS (5.2 ± 3.5 × 10^−7^) (14), the most successful genome-wide consensus sequencing approach to now.

**Table 2.** Number of mutants detected in treated TK6 with 6 bp trimmed from the distal regions.

| Treatment | Consensus bases<br>($\times 10^8$) | Total<br>mutants | Total-MF<br>($\times 10^{-6}$) | SNV | SNV-MF<br>($\times 10^{-6}$) | Deletion | Insertion | Indel-MF<br>($\times 10^{-6}$) |
| --- | --- | --- | --- | --- | --- | --- | --- | --- |
| Control | 2.79 | 114 | 0.41 | 111 | 0.40 | 2 | 1 | 0.01 |
| MMS | 2.81 | 149 | 0.53 | 144 | 0.51 | 4 | 1 | 0.01 |
| ENU | 2.11 | 229 | 1.08 | 225 | 1.07 | 4 | 0 | 0.02 |

### Low-level artifacts in the background mutant profile

Mutant profile of the 6 bp-trimmed control data revealed a relatively higher mutant frequency in CG base pair (3.26 × 10^−7^) compared to TA base pair (7.16 × 10^−8^) and CG > GC transversion (41.44%) was the most predominant base substitution type followed by CG > AT transversion (26.13%) and CG > TA transition (14.41%) (Figure 4). Compared with other well-established background mutational spectra (29,30), the CG > GC transversion, which is supposed to account for a relatively low proportion, was dominant in our data, suggesting some artifacts, at least in the CG base pairs, are still included even after correcting the bias in distal regions.

**Figure 4.**
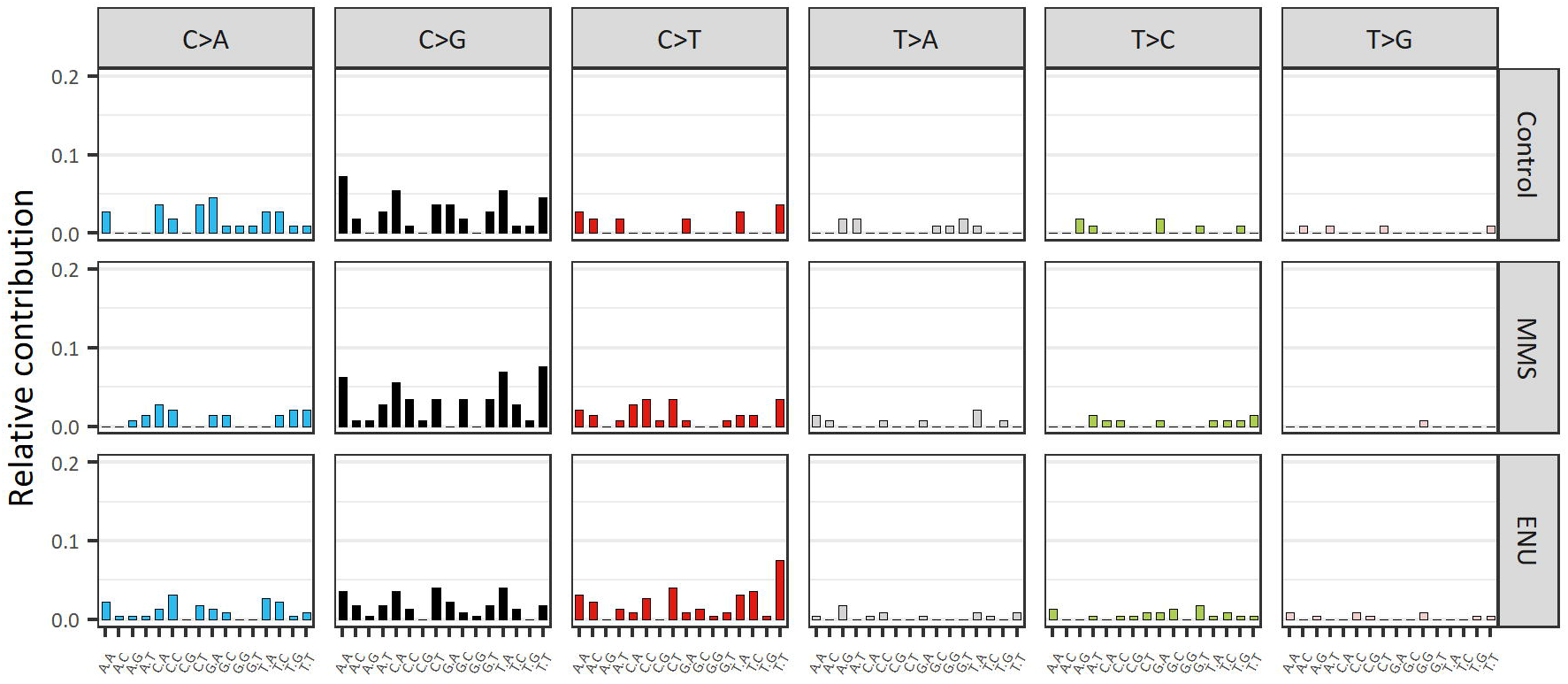
Profiles of mutants detected in terminal 6 bp-trimmed data. After trimming 6 base pairs from the distal regions, 111, 144 and 225 SNVs from the control, MMS-treated and ENU-treated data were obtained for the mutation spectra analysis, respectively.

During the fragmentation process, besides double strand breakage, nicks can also be produced, which are then subjected to the same error-prone end-repair process as the terminal overhangs (4). Notably, the mutation spectrum of the control data (Figure 4) showed a similar pattern as that of artifacts in distal regions (Figure 3) (cosine similarity = 0.827), suggesting the end-repair artifacts may also happen in the interior regions, even though with a much lower frequency compared to that of the distal regions.

To further confirm that the artifacts in the background mutant profile were the errors introduced during library preparations rather than the bias arising from PECC-Seq analysis, we applied PECC-Seq to the genomic library (DMSO-treated TK6) prepared with the enzymatic fragmentation method. From the enzymatic fragmented library, a total of 401 mutants (379 SNVs, 13 deletions and 9 insertions) and 1.78 × 10^7^ consensus bases were obtained, resulting in an error frequency of 2.25 × 10^−5^. As expected, different patterns of mutant distribution and base substitution types were observed. In contrast to the data from mechanical fragmented libraries, only the first base pair in the terminus exhibited a significantly higher error frequency compared with the interior regions (4.08 × 10^−4^ of 1^st^ bp to the terminus vs. 1.72 × 10^−5^ of 1 bp-trimmed data, *p* < 0.01, Figure S2). In addition, the most common substitution types observed in the enzymatic fragmented library were CG > TA transition, TA > CG transition and TA > AT transversion (Figure 5). Noteworthy, CG > GC transversion, the most predominant mutant type in mechanical fragmented libraries, was rare with only 1 among 1.78 × 10^7^ consensus bases in the enzymatic fragmented library. The different mutation spectra observed in mechanical fragmented (Figure 4) and enzymatic fragmented (Figure 5) libraries (cosine similarity = 0.168) further supported the hypothesis that the bias observed in the background mutant profile was more like the low-level artifacts produced during mechanical library preparations and may originate from the end-repair process, but should not be the bias arising from PECC-Seq analysis.

**Figure 5.**
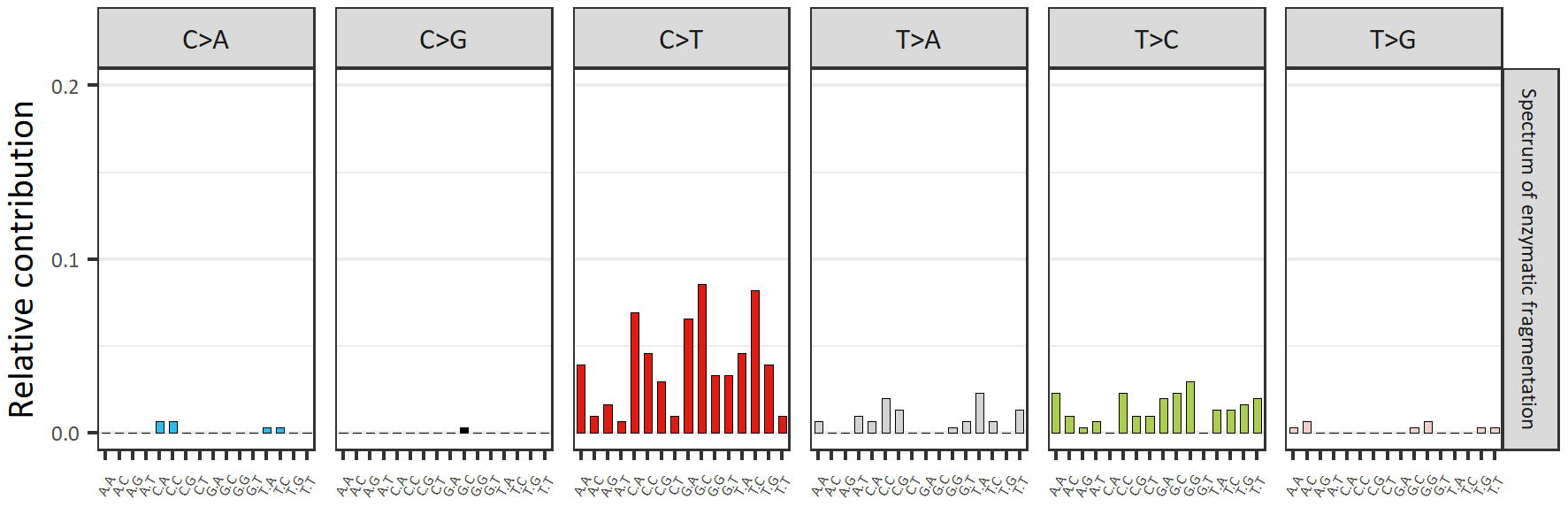
Profile of mutants detected in the enzymatic fragmented library of the control TK6. Mutants within 1 bp to the terminus were considered as end-repair artifacts and trimmed off. Total 304 SNVs were obtained and subjected to the spectrum analysis.

### Performance of PECC-Seq in detecting ENU- and MMS-specific mutations

In order to assess the ability of PECC-Seq in detecting low-frequency chemical-induced mutations at the whole genome scale, the mutant data from MMS-treated and ENU-treated TK6 were compared with the control data (Figure 4).

For MMS group, bases at the position of 7 bp to the terminus not only exhibited a much higher mutant frequency compared to the interior regions (Figure 2) and shared a similar mutant pattern as artifacts in distal regions (12.5% CG > AT, 75% CG > GC and 12.5% CG > TA), indicating that mutants exist at this site may still be the end-repair artifacts rather than true mutations. Thus, we analyzed the MMS data with the terminal 7 bp trimmed. As shown in Figure 6, the expected CG > TA transition, which is supposed to be specific to MMS-treatment (31,32), exhibited a relatively higher increase (1.86-fold) compared with other base substitution types.

**Figure 6.**
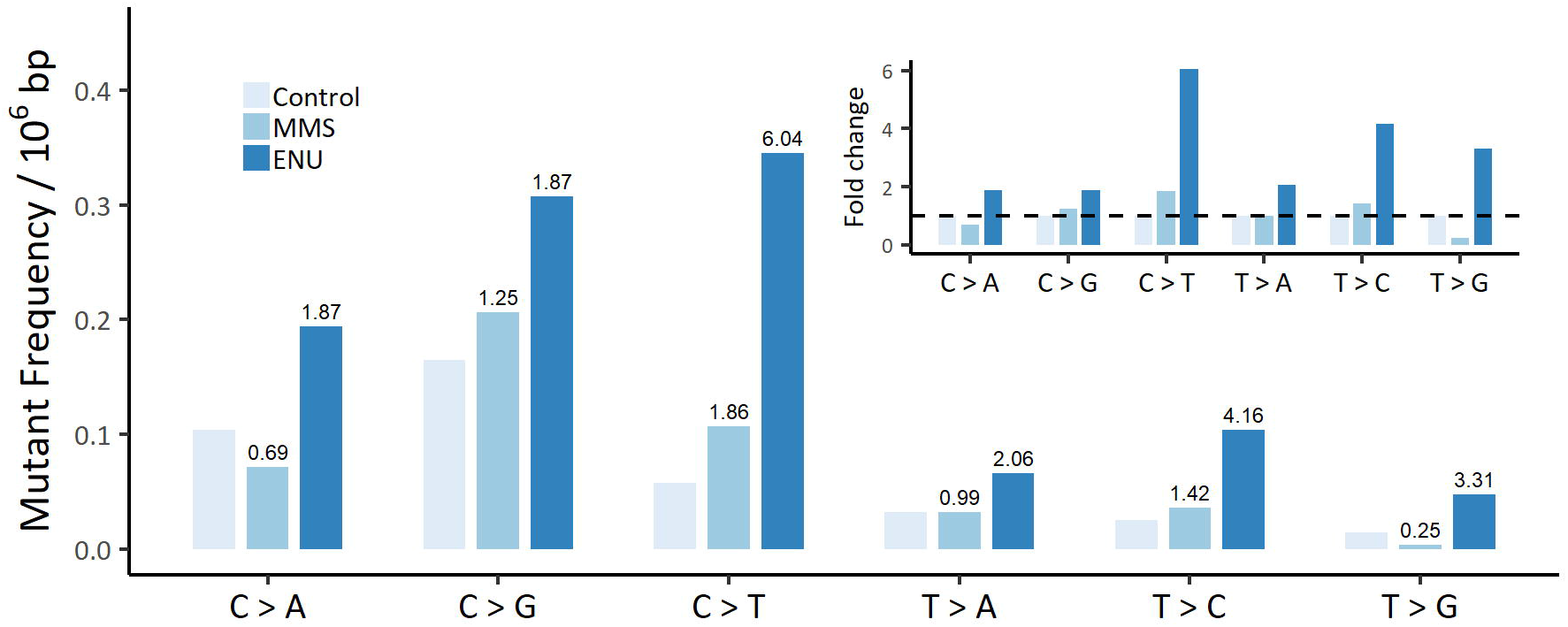
Mutant frequencies in MMS- and ENU-treated TK6 compared with the control data. The main figure shows the 6 base substitution types-specific mutant frequencies in the control and treated TK6. Number on the top of the bars indicates the relative fold changes in the mutant frequencies of each substitution type compared with the corresponding control data as displayed in the insert. For MMS-treated TK6, terminal 7 bp-trimmed data were used for comparison.

The ENU-treated TK6 exhibited a mutant frequency of 1.07 × 10^−6^ for SNVs with a more than 2-fold increase compared with the control. In TA base pairs, which are susceptible to ENU treatment (33,34), a 4.16-fold increase was observed in TA > CG transition, followed by TA > GC transversion (3.31-fold increase) and TA > AT transversion (2.06-fold increase). Unexpectedly, CG > TA transition also exhibited a 6.04-fold increase compared to the corresponding control data.

## DISCUSSION

Molecular consensus sequencing strategy has provided an optimal solution so far for enhancing the accuracy of NGS. Based on the principle of utilizing redundant copies derived from individual DNA molecules for error correction, this strategy usually requires a relatively higher sequencing depth, which makes it less cost-efficient when applied to large genomic targets (e.g., human genome). To seek a balance between accuracy, cost and sequencing yield, here, we present PECC-Seq, which improves the consensus-making efficiency by capitalizing the minimum number of same molecules-derived copies for error correction as demonstrated with the whole genome sequencing of TK6.

In contrast to other consensus sequencing strategies, PECC-Seq employs a PCR-free library for consensus sequencing, which minimizes the raw depth and the resultant cost for sequencing and avoids artifacts introduced by PCR amplification. Generally, PCR amplification is applied during library preparations to obtain sufficient copies for consensus sequencing. In PECC-Seq, we utilized the overlap in the paired-end reads and reverse complementary fragments arising from the duplex DNA libraries to obtain 4 reads for duplex consensus sequencing without using PCR amplification (Figure 1). Theoretically, the error frequency of the consensus sequencing approach can be roughly calculated as (the error rate of the sequencer × 1/3) ^number of reads to make consensus sequences^ × 3 (at a given site, the base can mutate to any one of other three types of bases) without considering the preference for different types of substitution errors. With 4 reads to create consensus sequences, the theoretical accuracy for PECC-Seq can reach a level lower than 10^−8^ with the sequencing error rate of 10^−3^ to 10^−2^ (Table S1). In addition, as the 2 pairs of paired-end reads are from the duplex templates, single strand damages-induced errors can be efficiently removed (4,15). Compared with the non-uniformity of PCR amplification-based consensus sequencing approaches (19,20), as the 4 copies are naturally generated from each library fragment, a uniform coverage of the genome can be acquired in PECC-Seq theoretically. Thus, by utilizing the minimum redundancy for consensus sequencing, relatively high accuracy and consensus-making efficiency and a uniform coverage can be achieved with decreased raw sequencing depth in PECC-Seq. Furthermore, PCR itself is an error-prone process and can introduce artifacts to the sequencing library. The errors that are introduced in the first several cycles of PCR are sometimes difficult to remove, especially in some single-strand consensus sequencing methods (4,11). With a PCR-free library, these hard-to-correct PCR artifacts can be avoided in PECC-Seq.

BotSeqS, which employed a dilution procedure to increase the consensus-making efficiency, is one of the very few genome-wide consensus sequencing approaches and has been successfully applied in detecting somatic mutations in human tissues (14). Compared with this PCR amplification-based duplex consensus sequencing approach, PECC-Seq exhibited a similar level of accuracy (4.1 × 10^−7^ in PECC-Seq vs. 5.2 ± 3.5 × 10^−7^ in BotSeqS). Noteworthy, with the 2 ×150 bp, 40 × sequencing depth data, and a recovery efficiency (i.e., the proportion of complementary reads in total paired-end reads) of approximately 1.2%, 2.57 × 10^8^ consensus bases and approximately 8% genome coverage on average could be acquired (Table S2). While in BotSeq libraries, only 1.03 × 10^7^ consensus bases and 0.4% genome coverage on average could be obtained with the 2 × 100 bp, 30 × sequencing depth data ((14), supplementary data). Thus, with a comparable high accuracy, PECC-Seq displayed a higher data yield with nearly a 20-fold increase compared with BotSeqS, revealing a much better cost-yield efficiency in genome-wide low-frequency mutation detection. In the standard sequencing procedure for HiSeqX, dsDNA library was denatured to single-strand fragments first and then diluted for sequencing. The dilution procedure can cause loss of the complementary strands derived from the same duplex templates. Thus, in the current data, only approximately 1.2% of the reads were from complementary strands, revealing an unsatisfied performance in recovering the complementary strands. To increase the proportion of same templates-derived complementary strands in the denatured and diluted sequencing libraries, reducing the input amount of library for denature and decreasing the subsequent dilution factor can be a simple but feasible solution. Therefore, by improving recovery efficiency of the complementary reads, there is still scope for further improvement on the data yield efficiency of PECC-Seq, which could enhance its performance in genome-wide mutation detection.

Random shearing process during fragmentation produces dsDNA fragments with blunt ends as well as large amounts of 3’ and 5’ overhangs, which require further end repair to generate uniform blunt ends before A-tailing and adaptor ligation (35). Usually, the 3’ overhangs are trimmed by 3’ to 5’ exonuclease, while the 5’ overhangs are fixed by DNA polymerases (36). Considering the error propensity of DNA polymerases, misincorporations may occur when processing repair of the overhangs, especially when nucleotide damages exist. Thus, end repair process is error-prone and may introduce errors to the terminal regions of the library fragments. In mechanical fragmented library, we observed that the distal region (6 bp to the terminus) is more susceptible to artifacts (Figure 2), revealing an end-repair error frequency of approximately 10^−5^ level. These end-repair artifacts feature a strong preference to CG base pairs in trinucleotide context of 5’-NpCpT-3’ or 5’-NpCpA-3’ and CG > GC transversion accounts for the most common substitution type. In contrast, in enzymatic fragmented library, only the 1^st^ bp in the terminal region showed a significantly increased error rate (Figure S2). This is consistent to the mechanisms of some enzymatic fragmentation methods in which short overhangs are produced (37). In addition, the error preference on 5’-NpCpT-3’ or 5’-NpCpA-3’ was not observed in terminal regions of the enzymatic fragmented library. These data suggested that library preparation procedures of different mechanisms may produce artifacts with different patterns of distribution and substitution types, and the characteristic artifacts observed in the context of 5’-NpCpT-3’ or 5’-NpCpA-3’ are the products specific to mechanical fragmentation-based library preparations. By removing the end-repair artifacts in the distal regions, the background accuracy of PECC-Seq was improved by about 2-fold. However, as suggested in our data, low-level end-repair artifacts can also occur in interior regions, which may mask the true mutations. Thus, characterizing the origins and patterns of such mechanical fragmented process-related artifacts may help to reduce and correct these errors by using modified experimental conditions or computational methods and further improve the sequencing accuracy.

High energy applied during ultrasonic shearing process can oxidize bases and introduce artifacts. CG > AT transversion is the most common substitution error observed in ultrasonic shearing libraries, which is generated by the misincorporation of adenine to oxidized guanine (i.e., 7,8-dihydro-8-oxoguanine, 8-oxoG) (28,38). In contrast, CG > GC transversion rather than CG > AT transversion exhibited the predominant artifacts in our data. Compared to other damages-related base substitution types (e.g., CG > AT transversion induced by 8-oxoG, CG > TA transition induced by deamination of cytosine), the mechanisms of CG > GC transversion are still not well-understood. Under oxidative pressure, in guanine, besides the typical 8-oxoG, various other oxidative products including 8-nitroguanine (8-NO2-G), 2-aminoimidazolone (Iz), 5-guanidino-4-nitroimidazole (NI), dehydroguanidinohydantoin (Ghox), and guanidinohydantoin (Gh), etc. are produced (39,40). A few studies have reported that Iz can induce mismatches mainly with Iz:G, which is then fixed into CG > GC transversion (39,41,42). One study has observed that Iz is the main product formed in single-stranded DNA (ssDNA) (40). Thus, we speculate that in the overhangs and nick-existing regions, the guanine in the produced ssDNA may tend to be oxidized into Iz rather than 8-oxoG, which results in more CG > GC transversion; while in double-stranded regions, the guanine is mainly oxidized into 8-oxoG, and such single strand damages-induced artifacts can be removed as complementary strands are utilized for error correction in PECC-Seq (Figure 7A). This hypothesis may explain the higher proportion of CG > GC transversion than CG > AT transversion in our data. Figure 7B shows a possible explanation for the difference in substitution types of artifacts observed in our data and other researches as different consensus sequencing strategies were applied. In some single-strand consensus sequencing methods, the single strand damages can be fixed into hard-to-correct errors after several cycles of PCR and then manifest as CG > AT transversion in the consensus sequences. Similarly to PECC-Seq, in DupSeq, with the information from complementary strands, the oxidative damages-induced artifacts can be efficiently removed, revealing suppressed CG > AT transversion errors (15). Noteworthy, in a previous study, we observed a relatively high proportion of CG > GC transversion in the data processed with DupSeq (43), indicating the occurrence of similar damages-induced artifacts. Our hypothesis may provide a possible explanation for the CG > GC transversion in this DupSeq data also. In addition, during library preparations, as a prolonged fragmentation procedure was applied in PECC-Seq, the generated 8-oxoG may be further oxidized to Iz (42), which may lead to fewer CG > AT transversions but more CG > GC transversions. However, the generation of Iz during mechanical fragmentation and its role in the CG > GC bias require further investigation.

**Figure 7.**
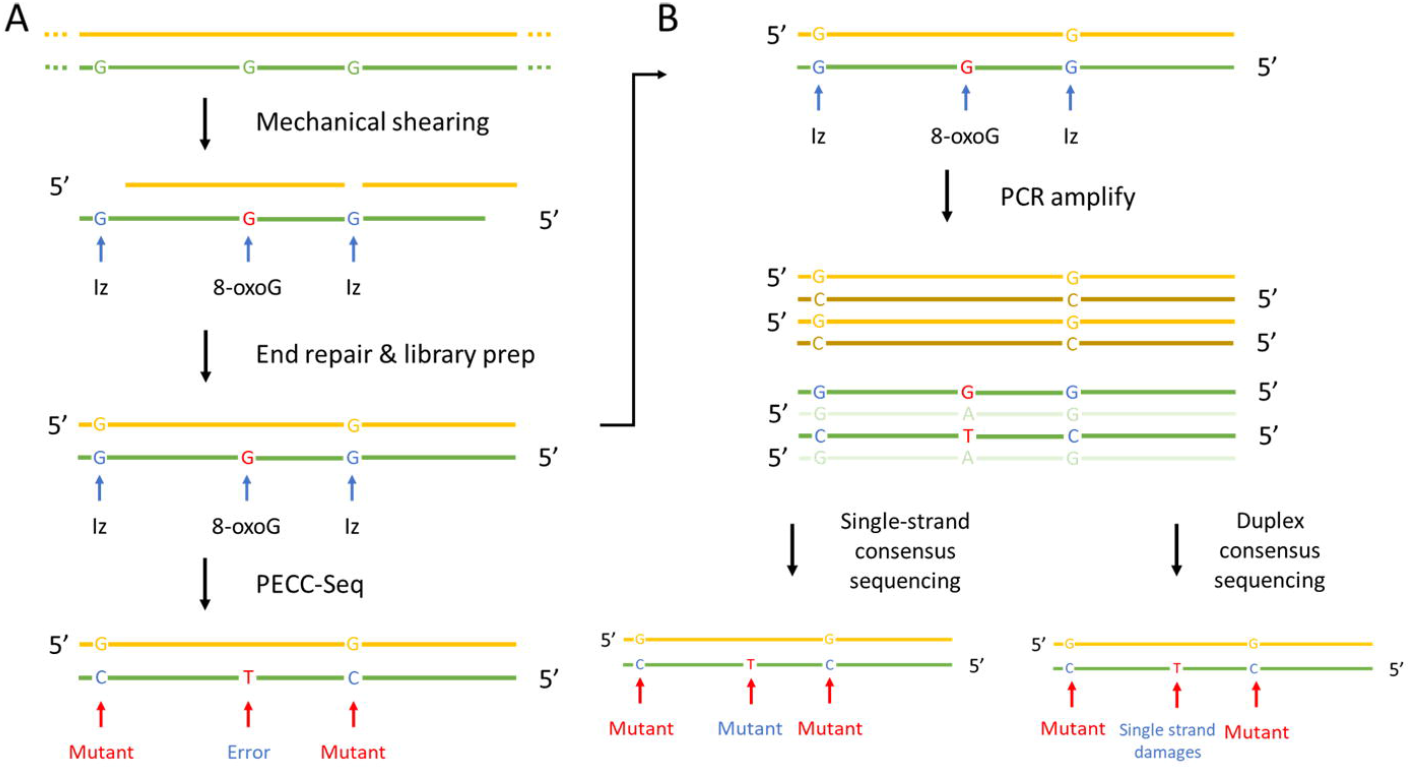
Possible mechanisms for the formation of end-repair artifacts. (**A**) A possible mechanism for the formation of the excess CG > GC transversion. We hypothesize that the guanine in ssDNA (in overhangs or nicked sites) may be oxidized to Iz, resulting in CG > GC transversion. In the double-stranded region, the guanine may be mainly oxidized into 8-oxoG, and such single strand damages-induced artifacts can be removed in PECC-Seq. (**B**) Possible explanations for the different spectra of mechanical fragmentation process-induced artifacts observed in our data and previous researches. After several cycles of PCR, the oxidative damages can be fixed into stable mutations. Based on the principles, single strand damages-induced artifacts can be efficiently removed in strategies that utilizing the information from the complementray strands, e.g., Duplex Sequencing, while these artifacts are difficult to be eliminated in some single-strand consensus sequencing methods.

We further evaluated the performance of PECC-Seq in detecting chemical-induced mutations. MMS and ENU are two commonly used mutagens in the gene mutation assays, both of which mainly induce single nucleotide substitutions (31–34). After treatment, the *TK* gene mutation frequencies exhibited significant increases in MMS-treated and ENU-treated TK6 as shown in the *TK* gene mutation assay (Table S3), indicating that the treatments indeed induced mutations in TK6. Since the chemical-induced effects may be masked by the low-level artifacts (especially artifacts of CG > GC transversion), we compared the mutant frequencies of chemical-specific base substitution types instead of the total mutant frequencies. In both data, increases in the chemical-specific base substitution types were detected. In MMS data, only a slight increase in MMS treatment-specific CG > TA transition (31,32) was observed; while in ENU data, the induced mutant frequencies in TA base pair exhibited relatively higher levels of increases with approximately 3-fold changes. These were consistent with the results of *TK* gene mutation assay as ENU induced a higher mutation frequency in the *TK* gene compared with MMS (4.90 × 10^−5^ for ENU-treated TK6 vs. 1.93 × 10^−5^ for MMS-treated TK6). However, the mutant frequencies (per base pair) obtained from PECC-Seq were higher than those expected from *TK* gene mutation frequencies (per gene) by dividing the length of *TK* gene (approximately 700 bp in length of the amino acid coding region (44)) and with a rough assumption of 1/2 to 1/3 mutations were silent (i.e., 1.07 - 1.43 × 10^−9^ for the control, 4.14 - 5.51 × 10^−8^ for MMS-treated and 1.05 - 1.40 × 10^−7^ for ENU-treated TK6). Similarly, the induced increased fold changes in mutant frequencies observed in sequencing data were lower than that of *TK* gene mutation assay (4.90 × 10^−5^ for ENU-treated and 1.93 × 10^−5^ for MMS-treated TK6 vs. 0.05 × 10^−5^ for the control), suggesting the residual artifacts may interfere with the performance of PECC-Seq in detecting chemical-induced mutations. Noteworthy, in ENU data, an abnormally significant increase in CG > TA transition was observed. ENU can react with DNA and form various adducts with both guanine and thymidine including N-alkylpurines (N7G), O^6^-alkylG, O^4^-alkylT and O^4^-alkylT, etc. (45). As the O^6^-alkylG can be efficiently repaired in mammalian cells by alkylguanine DNA transferases, ENU mainly induces mutations at TA base pair rather than CG base pair (45). As mentioned before, mechanical shearing process may produce oxidative damages on guanines, we speculate that these oxidative damages may have an interaction with the ENU-induced damages on guanines and affect the efficient repair of alkylated guanines, which lead to an increase in CG > TA transition. These data indicate the ability of PECC-Seq in detecting mutations with frequencies as low as mid-10^−7^, which is adequate for the applications in the fields of cancer research and clinical liquid biopsy, etc. (10). To detect ultralow-frequency mutations (e.g., chemical-induced mutations) with PECC-Seq, further reduction on the low-level residual artifacts is required. As these low-level artifacts are difficult to remove solely by consensus sequencing-based error correction, optimization of the experimental conditions (e.g., using milder mechanical shearing procedures, adding antioxidants and using high-fidelity end-repair enzymes) to reduce the damages and artifacts introduced during library preparation are needed. In the present work, we performed the prolonged fragmentation to obtain shorter DNA fragments in order to get overlaps in paired-end reads. With the advent of some sequencers with longer read length (e.g., MiSeq), PECC-Seq can be applied to library with longer DNA fragments. This means a milder mechanical fragmentation procedure can be applied, which may reduce the mechanical fragmentation-related artifacts. Furthermore, with longer read length, the variants introduced by mapping errors may also be further suppressed.

None of the approaches, however, is optimal for all the situations. In PECC-Seq, as the shear points are used as the endogenous barcode, one challenge is “tag clashes” when applied to deep sequencing (4,46). With the increase of sequencing depth, the probability of two individual templates by chance to have same mapping coordinates will increase. Wrongly pairing the two reads from different templates may exclude the true mutations and lead to false negative (4). We had also applied PECC-Seq to the analysis of mitochondrial DNA (which was an original target) from the control TK6. As a mtDNA-enriched DNA extraction kit was applied, the average sequencing depth of mtDNA was over 15,000 ×. With the same analysis pipeline, the resulting mutant frequency of mtDNA was lower than 4.21 × 10^−7^ (0 in 2.37 × 10^6^ consensus bases), which was much lower than the established background mtDNA mutation frequency of approximately 10^−5^ (15). This suggested that a false negative tendency was introduced by PECC-Seq when applied to small genomic targets with a high sequencing depth. To solve this problem in the scenario of deep sequencing for rare mutations, employing exogenous tags in combination to increase the diversity of barcodes is a possible solution. In PECC-Seq, conventional exogenous barcodes may be difficult to be incorporated. In order to identify templates arising from DNA duplex, the two complementary strands must have identical barcodes, while the barcodes are always different for complementary fragments in conventional tagging systems designed for single-strand consensus sequencing. Furthermore, as PECC-Seq is based on a PCR-free library preparation, PCR amplification-based tagging methods are not applicable. The Duplex Tag, which is employed in DupSeq for duplex consensus sequencing, may be an ideal exogenous barcode system for PECC-Seq (15), as the duplex tags can provide identical barcode information for a pair of duplex strands and can be tagged to DNA fragments with a ligation-based tagging method. Besides using exogenous barcodes, another simple solution is multiplexing the same samples that are labelled with different index sequences. By utilizing the index system, the total sequencing depth for a single sample can be multiplied several folds when applying PECC-Seq.

In conclusion, PECC-Seq exhibits a fairly high accuracy and a relatively good cost-yield efficiency in detecting low-frequency mutations at the whole-genome scale, which has potential applications in such fields as clinical liquid biopsy, cancer research and as an alternative for the gene mutation assay in future. In addition, with the accuracy of PECC-Seq, we demonstrated the end-repair artifacts that are introduced during mechanical fragmentation-based library preparations, which are prominent at the terminal 6 bp of the library fragments in trinucleotide context of 5’-NpCpA-3’ or 5’-NpCpT-3’. By characterizing and reducing such artifacts, the accuracy of consensus sequencing approaches, not only limit to PECC-Seq, may be further improved. Based on the results, we recommend to trim error-prone bases in the terminal regions of the sequenced fragments (bases within 6 to 7 bp to the terminus for mechanical fragmented libraries) to reduce end-repair artifacts in routine NGS data analysis.

## ACKNOWLEDGEMENTS

We appreciate Dr. Rajaguru Palanisamy for his critical reading and suggestions on our manuscript.

## REFERENCES

1. Kennedy, S.R., Schmitt, M.W., Fox, E.J., Kohrn, B.F., Salk, J.J., Ahn, E.H., Prindle, M.J., Kuong, K.J., Shen, J.C., Risques, R.A. et al. (2014) Detecting ultralow-frequency mutations by Duplex Sequencing. Nat Protoc, 9, 2586–2606.

2. Fox, E.J., Reid-Bayliss, K.S., Emond, M.J. and Loeb, L.A. (2014) Accuracy of Next Generation Sequencing Platforms. Next Gener Seq Appl, 1.

3. Shendure, J. and Ji, H. (2008) Next-generation DNA sequencing. Nat Biotechnol, 26, 1135–1145.

4. Salk, J.J., Schmitt, M.W. and Loeb, L.A. (2018) Enhancing the accuracy of next-generation sequencing for detecting rare and subclonal mutations. Nat Rev Genet, 19, 269–285.

5. Beckman, R.A. and Loeb, L.A. (2017) Evolutionary dynamics and significance of multiple subclonal mutations in cancer. DNA Repair (Amst), 56, 7–15.

6. Schmitt, M.W., Loeb, L.A. and Salk, J.J. (2016) The influence of subclonal resistance mutations on targeted cancer therapy. Nat Rev Clin Oncol, 13, 335–347.

7. Besser, J., Carleton, H.A., Gerner-Smidt, P., Lindsey, R.L. and Trees, E. (2018) Next-generation sequencing technologies and their application to the study and control of bacterial infections. Clin Microbiol Infect, 24, 335–341.

8. Yang, Y., Xie, B. and Yan, J. (2014) Application of next-generation sequencing technology in forensic science. Genomics Proteomics Bioinformatics, 12, 190–197.

9. Bragg, L. and Tyson, G.W. (2014) Metagenomics using next-generation sequencing. Methods Mol Biol, 1096, 183–201.

10. Vasan, N., Baselga, J. and Hyman, D.M. (2019) A view on drug resistance in cancer. Nature, 575, 299–309.

11. Sloan, D.B., Broz, A.K., Sharbrough, J. and Wu, Z. (2018) Detecting Rare Mutations and DNA Damage with Sequencing-Based Methods. Trends Biotechnol, 36, 729–740.

12. Kinde, I., Wu, J., Papadopoulos, N., Kinzler, K.W. and Vogelstein, B. (2011) Detection and quantification of rare mutations with massively parallel sequencing. Proc Natl Acad Sci U S A, 108, 9530–9535.

13. Lou, D.I., Hussmann, J.A., McBee, R.M., Acevedo, A., Andino, R., Press, W.H. and Sawyer, S.L. (2013) High-throughput DNA sequencing errors are reduced by orders of magnitude using circle sequencing. Proc Natl Acad Sci U S A, 110, 19872–19877.

14. Hoang, M.L., Kinde, I., Tomasetti, C., McMahon, K.W., Rosenquist, T.A., Grollman, A.P., Kinzler, K.W., Vogelstein, B. and Papadopoulos, N. (2016) Genome-wide quantification of rare somatic mutations in normal human tissues using massively parallel sequencing. Proc Natl Acad Sci U S A, 113, 9846–9851.

15. Schmitt, M.W., Kennedy, S.R., Salk, J.J., Fox, E.J., Hiatt, J.B. and Loeb, L.A. (2012) Detection of ultra-rare mutations by next-generation sequencing. Proc Natl Acad Sci U S A, 109, 14508–14513.

16. Matsumura, S., Sato, H., Otsubo, Y., Tasaki, J., Ikeda, N. and Morita, O. (2019) Genome-wide somatic mutation analysis via Hawk-Seq reveals mutation profiles associated with chemical mutagens. Arch Toxicol, 93, 2689–2701.

17. Li, C., Chng, K.R., Boey, E.J., Ng, A.H., Wilm, A. and Nagarajan, N. (2016) INC-Seq: accurate single molecule reads using nanopore sequencing. Gigascience, 5, 34.

18. Wenger, A.M., Peluso, P., Rowell, W.J., Chang, P.C., Hall, R.J., Concepcion, G.T., Ebler, J., Fungtammasan, A., Kolesnikov, A., Olson, N.D. et al. (2019) Accurate circular consensus long-read sequencing improves variant detection and assembly of a human genome. Nat Biotechnol, 37, 1155–1162.

19. Aird, D., Ross, M.G., Chen, W.S., Danielsson, M., Fennell, T., Russ, C., Jaffe, D.B., Nusbaum, C. and Gnirke, A. (2011) Analyzing and minimizing PCR amplification bias in Illumina sequencing libraries. Genome Biol, 12, R18.

20. Oyola, S.O., Otto, T.D., Gu, Y., Maslen, G., Manske, M., Campino, S., Turner, D.J., Macinnis, B., Kwiatkowski, D.P., Swerdlow, H.P. et al. (2012) Optimizing Illumina next-generation sequencing library preparation for extremely AT-biased genomes. BMC Genomics, 13, 1.

21. Honma, M. and Hayashi, M. (2011) Comparison of in vitro micronucleus and gene mutation assay results for p53-competent versus p53-deficient human lymphoblastoid cells. Environ Mol Mutagen, 52, 373–384.

22. OECD/OCDE. (2016) In Vitro Mammalian Cell Gene Mutation Tests Using the Thymidine Kinase Gene. Guidelines for the Testing of Chemicals, Section 4.

23. Bolger, A.M., Lohse, M. and Usadel, B. (2014) Trimmomatic: a flexible trimmer for Illumina sequence data. Bioinformatics, 30, 2114–2120.

24. OECD/OCDE. (2013) Transgenic Rodent Somatic and Germ Cell Gene Mutation Assays. Guidelines for the Testing of Chemicals, Section 4.

25. Roach, J.C., Glusman, G., Smit, A.F., Huff, C.D., Hubley, R., Shannon, P.T., Rowen, L., Pant, K.P., Goodman, N., Bamshad, M. et al. (2010) Analysis of genetic inheritance in a family quartet by whole-genome sequencing. Science, 328, 636–639.

26. Besenbacher, S., Liu, S., Izarzugaza, J.M., Grove, J., Belling, K., Bork-Jensen, J., Huang, S., Als, T.D., Li, S., Yadav, R. et al. (2015) Novel variation and de novo mutation rates in population-wide de novo assembled Danish trios. Nat Commun, 6, 5969.

27. Milholland, B., Dong, X., Zhang, L., Hao, X., Suh, Y. and Vijg, J. (2017) Differences between germline and somatic mutation rates in humans and mice. Nat Commun, 8, 15183.

28. Peng, Q. and Xu, C. (2019) Targeted Single Primer Enrichment Sequencing with Single End Duplex-UMI. Sci Rep, 9, 4810.

29. Kucab, J.E., Zou, X., Morganella, S., Joel, M., Nanda, A.S., Nagy, E., Gomez, C., Degasperi, A., Harris, R., Jackson, S.P. et al. (2019) A Compendium of Mutational Signatures of Environmental Agents. Cell, 177, 821–836.e816.

30. Behjati, S., Huch, M., van Boxtel, R., Karthaus, W., Wedge, D.C., Tamuri, A.U., Martincorena, I., Petljak, M., Alexandrov, L.B., Gundem, G. et al. (2014) Genome sequencing of normal cells reveals developmental lineages and mutational processes. Nature, 513, 422–425.

31. Op het Veld, C.W., Jansen, J., Zdzienicka, M.Z., Vrieling, H. and van Zeeland, A.A. (1998) Methyl methanesulfonate-induced hprt mutation spectra in the Chinese hamster cell line CHO9 and its xrcc1-deficient derivative EM-C11. Mutat Res, 398, 83–92.

32. Klungland, A., Laake, K., Hoff, E. and Seeberg, E. (1995) Spectrum of mutations induced by methyl and ethyl methanesulfonate at the hprt locus of normal and tag expressing Chinese hamster fibroblasts. Carcinogenesis, 16, 1281–1285.

33. Takahasi, K.R., Sakuraba, Y. and Gondo, Y. (2007) Mutational pattern and frequency of induced nucleotide changes in mouse ENU mutagenesis. BMC Mol Biol, 8, 52.

34. Barbaric, I., Wells, S., Russ, A. and Dear, T.N. (2007) Spectrum of ENU-induced mutations in phenotype-driven and gene-driven screens in the mouse. Environ Mol Mutagen, 48, 124–142.

35. Bronner, I.F. and Quail, M.A. (2019) Best Practices for Illumina Library Preparation. Curr Protoc Hum Genet, 102, e86.

36. Illumina. (2017) TruSeq DNA PCR-Free Reference Guide. https://support.illumina.com.cn/downloads/truseq-dna-pcr-free-reference-guide-1000000039279.html. Accessed 26 Nov 2019.

37. Knierim, E., Lucke, B., Schwarz, J.M., Schuelke, M. and Seelow, D. (2011) Systematic comparison of three methods for fragmentation of long-range PCR products for next generation sequencing. PLoS One, 6, e28240.

38. Costello, M., Pugh, T.J., Fennell, T.J., Stewart, C., Lichtenstein, L., Meldrim, J.C., Fostel, J.L., Friedrich, D.C., Perrin, D., Dionne, D. et al. (2013) Discovery and characterization of artifactual mutations in deep coverage targeted capture sequencing data due to oxidative DNA damage during sample preparation. Nucleic Acids Res, 41, e67.

39. Neeley, W.L., Delaney, J.C., Henderson, P.T. and Essigmann, J.M. (2004) In vivo bypass efficiencies and mutational signatures of the guanine oxidation products 2-aminoimidazolone and 5-guanidino-4-nitroimidazole. J Biol Chem, 279, 43568–43573.

40. Morikawa, M., Kino, K., Oyoshi, T., Suzuki, M., Kobayashi, T. and Miyazawa, H. (2014) Analysis of guanine oxidation products in double-stranded DNA and proposed guanine oxidation pathways in single-stranded, double-stranded or quadruplex DNA. Biomolecules, 4, 140–159.

41. Kino, K. and Sugiyama, H. (2001) Possible cause of G-C-->C-G transversion mutation by guanine oxidation product, imidazolone. Chem Biol, 8, 369–378.

42. Kino, K., Hirao-Suzuki, M., Morikawa, M., Sakaga, A. and Miyazawa, H. (2017) Generation, repair and replication of guanine oxidation products. Genes Environ, 39, 21.

43. Chawanthayatham, S., Valentine, C.C.3rd, Fedeles, B.I., Fox, E.J., Loeb, L.A., Levine, S.S., Slocum, S.L., Wogan, G.N., Croy, R.G. and Essigmann, J.M. (2017) Mutational spectra of aflatoxin B1 in vivo establish biomarkers of exposure for human hepatocellular carcinoma. Proc Natl Acad Sci U S A, 114, E3101–e3109.

44. Liber, H.L., Yandell, D.W. and Little, J.B. (1989) A comparison of mutation induction at the tk and hprt loci in human lymphoblastoid cells; quantitative differences are due to an additional class of mutations at the autosomal tk locus. Mutat Res, 216, 9–17.

45. Jenkins, G.J., Doak, S.H., Johnson, G.E., Quick, E., Waters, E.M. and Parry, J.M. (2005) Do dose response thresholds exist for genotoxic alkylating agents? Mutagenesis, 20, 389–398.

46. Schmitt, M.W., Fox, E.J. and Salk, J.J. (2014) Risks of double-counting in deep sequencing. Proc Natl Acad Sci U S A, 111, E1560.

