## Supplementary for "Detection of genome-wide low-frequency mutations with Paired-End and Complementary Consensus Sequencing (PECC-Seq) revealed end-repair derived artifacts as residual errors"

**SUPPLEMENTARY INFORMATION**

**Supplementary Materials and Methods**

**Sequencing data preprocessing**

The raw sequencing data were first subjected to read trimming with the Trimmomatic software (v0.39) (1). Since shortened library fragments were utilized in PECC-Seq, “adapter read-through” might occur, which could result in adapter contamination at the end of the reads. Thus, the “palindrome mode” was applied to ensure the efficient adapter removal (1). As the 5’ mapping coordinates of the reads served as endogenous barcodes to identify the complementary strands, no 5’-terminal trimming was applied to the reads. In addition, to retain maximum information of the reads, no quality filtering was performed at this step. After trimming, the paired-end reads were aligned to human reference genome hg38 with the Burrows-Wheeler aligner (BWA) followed by sorting with SAMtools (v1.9) (2).

As the accuracy of endogenous barcodes highly depends on the proper mapping of the reads, strict read filtering was applied after mapping. Read filtering was performed using SAMtools with the command as follows: $ samtools view -F 2316 -f 2 -q 60. After removing the non-mapping reads, and the reads with multiple mappings, only properly mapped paired-end reads (i.e., both reads of the read pairs are properly mapped) with mapping quality of 60 were retained.

**Extraction of consensus read groups**

Extraction of the consensus read groups was performed with the R software (v1.2.1335). The 5’ mapping coordinates of the paired-end reads were extracted, i.e., the 1-based leftmost position and the 1-based rightmost position of the paired-end reads. Paired-end reads with soft clipping at 5’ ends were discarded. Read pairs with same 5’ mapping coordinates were then picked up and grouped. Groups with more than two pairs of paired-end reads were discarded as they might contain optical duplicates or different templates-derived reads that have matched mapping coordinates by chance. As the fragments derived from the duplex templates should be reverse complementary, the mapping orientations of complementary fragments are opposite. Thus, the FLAG of the 4 reads (i.e., 2 pairs of paired-end reads) in a consensus read group should be 83 (= 1 + 2 + 16 + 64, the read is the first read (64) from the properly mapped paired-end reads (1 + 2) and is mapped to reference genome on the reverse strand (16)), 163 (= 1 + 2 + 32 + 128, the read is the second read (128) from the properly mapped paired-end reads (1 + 2) and its mate is mapped to reference genome on the reverse strand (32)), 99 (= 1 + 2 + 32 + 64, the read is the first read (64) from the properly mapped paired-end reads (1 + 2) and its mate is mapped to reference genome on the reverse strand (32)) and 147 (= 1 + 2 + 16 + 128, the read is the second read (128) from the properly mapped paired-end reads (1 + 2) and is mapped to reference genome on the reverse strand (16)). By filtering with the FLAG, groups with reads of same mapping orientations were excluded and only duplex DNA-derived complementary paired-end reads were retained. Further, consensus read groups that mapped to regions with ≥ 100 × depth (approximately 2-fold of average sequencing depth) were discarded as these regions may contain repetitive sequences, which could result in misalignment or wrong pairing of the complementary reads.

**Extraction of consensus bases**

The 4 sequences in the overlap region of the consensus read groups (main text, Fig. 1C) were extracted to form consensus sequences. At a given site, the 2 bases in the overlap of the pair-end reads were utilized to make the “paired-end consensus base” first. Mismatches between these two overlapped bases reflected the sequencing errors. Then the paired-end consensus bases were further subjected to form “complementary consensus bases”. Mismatches of the two complementary strands-derived paired-end consensus bases were considered as a mixture of sequencing errors, single-strand artifacts, and errors introduced by wrong grouping of reads arising from different templates. Such mismatches that resulted from sequencing errors and single-strand artifacts should be randomly distributed. The probability of two random errors happen in same reads was very low (*Poisson* test, *p* < 0.01), while mismatches introduced by wrong pairing two individual templates might have a higher probability to happen in the same consensus read groups. Thus, consensus read groups with more than 3 mismatches between paired-end consensus bases were discarded at this stage empirically. Finally, only positions with 4 identical bases were included for further mutation analysis.

**Mutation analysis**

The resulting consensus bases were then filtered and subjected to the mutation analysis. As newly generated mutations should only present in single cells, the candidate mutations were picked up with the following criteria: 1) variants happened solely in the consensus reads and not found in other reads of the variants site; and 2) variants were not found in data of other treatment groups. By utilizing information from other treatment groups, mutants derived from potential single nucleotide polymorphisms (SNPs) could be removed. Consensus reads with more than 2 mutants present in the same reads were discarded empirically. All candidate mutants were further confirmed with the IGV browser.

**Overview of PECC-Seq data processing**

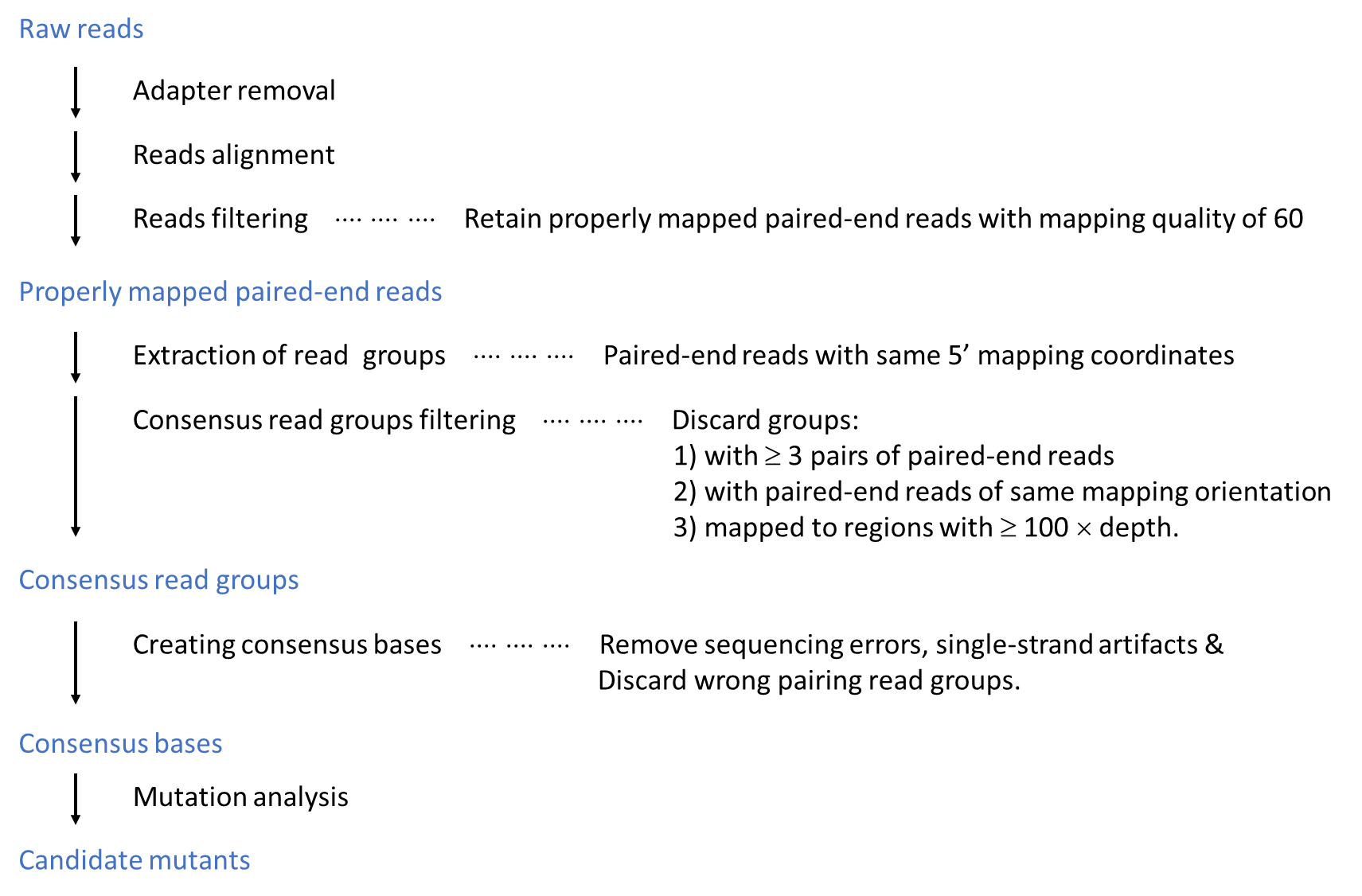

**Supplementary Results**

**Probability of “tag clashes” with current sequencing depth**

After the paired-end sequencing, total 9.42 × 10^8^, 9.10 × 10^8^ and 8.18 × 10^8^ raw sequencing reads were obtained from the control, MMS-treated and ENU-treated TK6 libraries, i.e., approximately 4.45 × 10^8^ paired-end reads on average with the current sequencing depth. The average insert sizes of the mapped paired-end reads were 186.0 ± 28.4, 174.6 ± 27.8 and 159.4 ± 27.0 (Mean ± SD) for the control, MMS-treated and ENU-treated data, respectively. With an assumption of a 20-bases variation in the fragment length, total number of available shear points could be roughly calculated as 3 × 10^9^ (approximate number of genomic coordinates) × 20 (number of variations in the fragment length), which equaled to 6 × 10^10^. The probability of 2 paired-end reads by chance to have same coordinates follows the *Poisson* distribution and was 2.7 × 10^-5^ (*Poisson* test) with a conservative estimation. Thus, the probability of the “tag clashes” scenario was under an acceptable level and endogenous barcodes can be utilized in the genome-wide sequencing strategy with the current sequencing depth.

**Table S1.** Error rates measured by different data processing methods.

| Data processing | | Control | | MMS | | ENU | |
| --- | --- | --- | --- | --- | --- | --- | --- |
|  |  | No. of bases (× 10^8^) | Error rate / Mutant frequency | No. of bases (× 10^8^) | Error rate / Mutant frequency | No. of bases (× 10^8^) | Error rate / Mutant frequency |
|  | Raw data^*^ | 11.73 | 4.69 × 10^-3^ | 11.86 | 4.06 × 10^-3^ | 9.08 | 3.88 × 10^-3^ |
|  | Paired-end consensus analysis^**^ | 5.75 | 1.40 × 10^-4^ | 5.83 | 1.10 × 10^-4^ | 4.46 | 1.43 × 10^-4^ |
|  | PECC-Seq  (raw data)^***^ | 2.83 | 0.76 × 10^-6^ | 2.88 | 1.14 × 10^-6^ | 2.22 | 2.85 × 10^-6^ |
|  | PECC-Seq  (6 bp-trimmed) | 2.79 | 0.41 × 10^-6^ | 2.81 | 0.53 × 10^-6^ | 2.11 | 1.08 × 10^-6^ |

* Raw data of extracted consensus read groups. Rough error rates were calculated as: Number of mismatches in paired-end consensus bases / Number of total bases.

** Data obtained from paired-end consensus bases. Rough error rates = Number of mismatches in the two complementary strands-derived paired-end consensus bases / Number of paired-end consensus bases.

*** Data of PECC-Seq (without trimming off the end-repair artifacts).

\# For consensus sequencing data, the number of bases indicates the number of consensus bases.

**Table S2.** Data yields and general performance of PECC-Seq.

| Treatment | PE reads (× 10^8^)^*^ | Consensus read groups (× 10^6^) | Recovery efficiency^**^ | Consensus bases (× 10^8^) | Coverage (× 10^8^ bp) | Coverage (%)^***^ | Depth |
| --- | --- | --- | --- | --- | --- | --- | --- |
| Control | 7.96 | 2.62 | 1.31% | 2.83 | 2.68 | 8.63% | 1.06 |
| MMS | 7.70 | 2.33 | 1.21% | 2.88 | 2.72 | 8.77% | 1.06 |
| ENU | 6.88 | 1.68 | 0.98% | 2.22 | 2.12 | 6.85% | 1.05 |

* Number of properly mapped paired-end reads after reads filtering.

** Recovery efficiency = Number of consensus read groups × 4 / Number of PE reads.

*** Coverage (%) = Coverage (bp) / whole genome size (bp).

**Table S3.** Mutation frequencies of the *TK* gene in treated TK6.

| Treatment | N-MF (× 10^-6^) | S-MF (× 10^-6^) | T-MF (× 10^-6^) |
| --- | --- | --- | --- |
| Control | 0.1 | 0.4 | 0.5 |
| MMS | 14.3^*^ | 5.0^*^ | 19.3^*^ |
| ENU | 38.3^*^ | 10.7^*^ | 49.0^*^ |

\# MF: mutation frequencies; N: Normally growing colony, S: Slowly growing colony, T: Total colony.

* *P* < 0.05 compared with the corresponding control data.

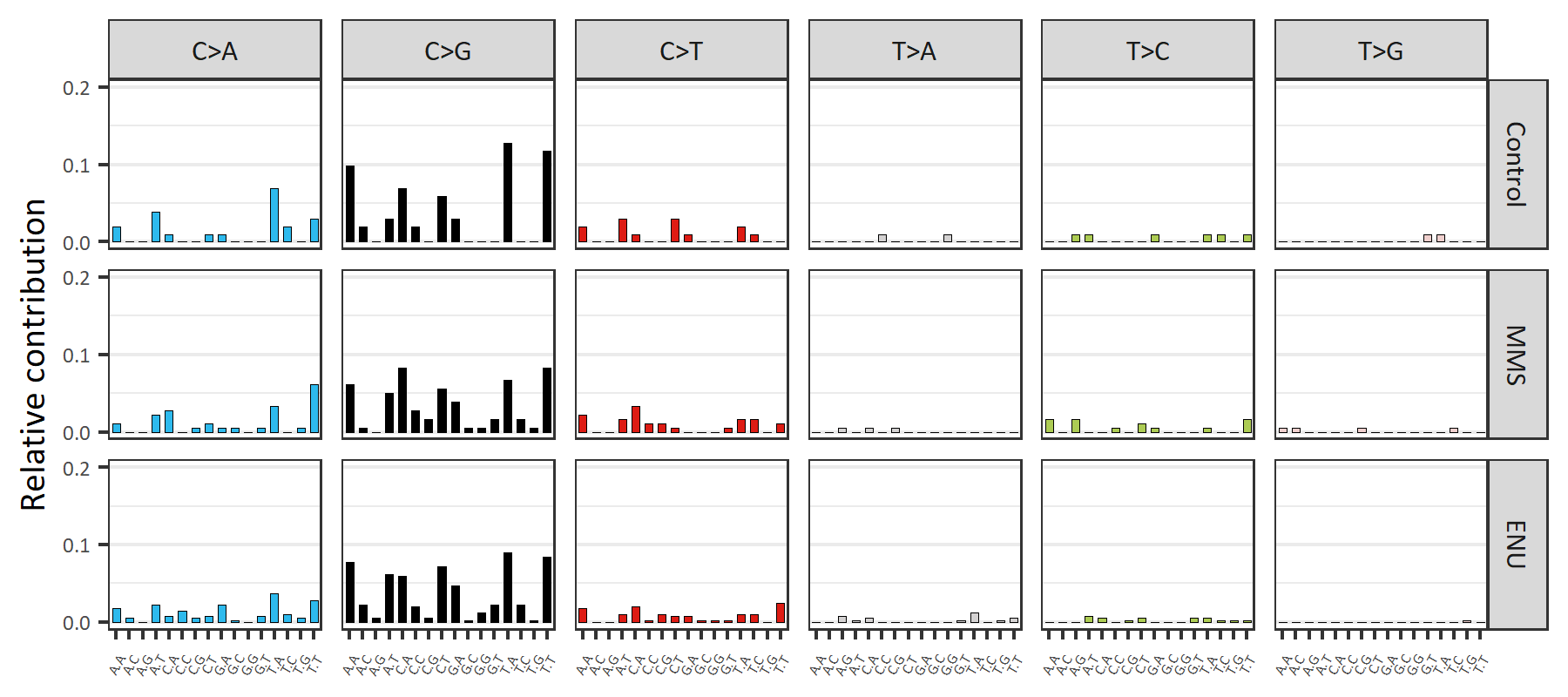

**Figure S1.** Profiles of mutants detected in the distal regions. The total 102, 180 and 403 SNVs detected in the distal regions of the control, MMS-treated and ENU-treated TK6 libraries were included for the mutation spectra analysis

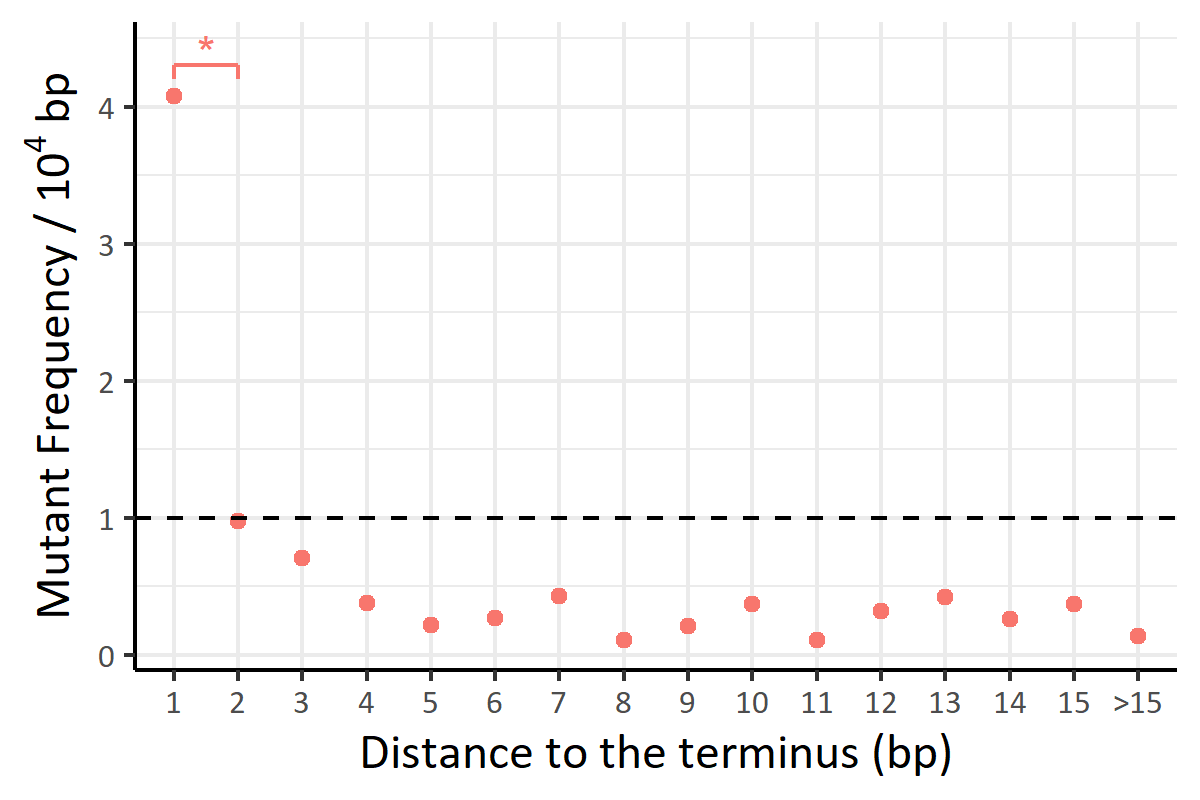

**Figure S2.** Mutant frequencies by positions in the enzymatic fragmented library. The terminus is defined as the beginning or end of the DNA templates (library fragments). * *p* < 0.05.

**References**

1.	Bolger, A.M., Lohse, M. and Usadel, B. (2014) Trimmomatic: a flexible trimmer for Illumina sequence data. *Bioinformatics*, **30**, 2114-2120.

2.	Li, H., Handsaker, B., Wysoker, A., Fennell, T., Ruan, J., Homer, N., Marth, G., Abecasis, G. and Durbin, R. (2009) The Sequence Alignment/Map format and SAMtools. *Bioinformatics*, **25**, 2078-2079.
